# α_2_-adrenergic signaling disrupts β cell BDNF-TrkB receptor tyrosine kinase signaling

**DOI:** 10.1101/400010

**Authors:** Michael A. Kalwat, Zhimin Huang, Derk D. Binns, Kathleen McGlynn, Melanie H. Cobb

**Affiliations:** Department of Pharmacology, UT Southwestern Medical Center, Dallas, TX 75235, USA; UT Southwestern Visiting Scholar

**Author notes:** Department of Endocrinology and Diabetes Center, First Affiliated Hospital of Sun Yat-sen University, 58 Zhongshan Er Road, Guangzhou, China, 510080 (Zhimin Huang). Correspondence: Michael Kalwat.

**Keywords:** cell signaling, pancreatic islet, extracellular-signal-regulated kinase (ERK), brain-derived neurotrophic factor (BDNF), BDNF/NT-3 growth factors receptor (TrkB), epinephrine, adrenergic receptor, diabetes

## Abstract

Adrenergic signaling is a well-known input into pancreatic islet function. Specifically, the insulin-secreting islet β cell expresses the G_i/o_-linked α_2_-adrenergic receptor, which upon activation suppresses insulin secretion. The use of adrenergic agonist epinephrine at micromolar doses may have supraphysiological effects. We found that pretreating β cells with micromolar concentrations of epinephrine differentially inhibited activation of receptor tyrosine kinases. We chose TrkB as an example because of its relative sensitivity to the effects of epinephrine and due to its potential regulatory role in the β cell. Our characterization of brain-derived neurotrophic factor (BDNF)-TrkB signaling in MIN6 β cells showed that TrkB is activated by BDNF as expected, leading to canonical TrkB autophosphorylation and subsequent downstream signaling, as well as chronic effects on β cell growth. Micromolar, but not nanomolar, concentrations of epinephrine blocked BDNF-induced TrkB autophosphorylation and downstream mitogen-activated protein kinase pathway activation, suggesting an inhibitory phenomenon at the receptor level. We determined epinephrine-mediated inhibition of TrkB activation to be G_i/o_-dependent using pertussis toxin, arguing against an off-target effect of high dose epinephrine. Published data suggested that inhibition of potassium channels or phosphoinositide-3-kinase signaling may abrogate the negative effects of epinephrine, however these did not rescue TrkB signaling in our experiments. Taken together, these results show that 1) TrkB kinase signaling occurs in β cells and 2) use of epinephrine in studies of insulin secretion requires careful consideration of concentration-dependent effects. BDNF-TrkB signaling in β cells may underlie pro-survival or growth signaling and warrants further study.

## 1 Introduction

Glucose homeostasis is largely controlled by the metered secretion of insulin from pancreatic islet β cells. β cells respond to elevated circulating glucose via coupling its metabolism to membrane depolarization, calcium (Ca^2+^) influx and insulin exocytosis (Kalwat and Cobb 2017). Secreted insulin suppresses liver gluconeogenesis and stimulates peripheral glucose uptake. Diabetes is a disease of hyperglycemia caused by deficient insulin production and action. In diabetes, β cells are either destroyed by the immune system (type 1 diabetes), or are unable to secrete sufficient insulin in response to stimulation (type 2 diabetes). In order to function properly, pancreatic β cells integrate a diverse array of inputs, including nutrients and hormones. To accomplish this, β cells utilize a variety of signaling mechanisms such as G-protein coupled receptors (GPCRs) (Holst 2007, Straub and Sharp 2012) and receptor tyrosine kinases (RTKs) (Kulkarni et al. 1999, Kulkarni et al. 2002, Song et al. 2016). Reported cross-talk between GLP1R and EGFR in islet β cells lends support to the idea of more general GPCR-RTK signaling interactions in β cells (Fusco et al. 2017).

Extracellular regulated kinase 1/2 (ERK1/2) is activated by insulin secretagogues (e.g. glucose, amino acids) and blunted by inhibitors of secretion (e.g. epinephrine), and is therefore frequently used as a proxy for β cell responsiveness (Longuet et al. 2005, Jaques et al. 2008, Goehring et al. 2011). ERK1/2 activation has long been recognized for its role in β cell growth and insulin gene expression (Hugl et al. 1998, Briaud et al. 2003, Khoo et al. 2003, Lawrence et al. 2008). Recently, acute ERK2 activity was demonstrated to be critical for the first-phase of insulin secretion (Leduc et al. 2017). Our interest in the pathways leading to ERK1/2 activation and the inhibitory functions of epinephrine in β cells led us to test the impact of epinephrine on RTK signaling to ERK1/2. Epinephrine has different effects on the ERK1/2 pathway depending on cell type and receptors expressed. In β cells, epinephrine activates α_2_-adrenergic receptors and inhibits insulin secretion as well as glucose-stimulated ERK1/2 activation (Peterhoff et al. 2003, Gibson et al. 2006).

We discovered that epinephrine suppressed RTK signaling in a concentration dependent manner and with varying potency depending on the RTK. Activation of α_2_-adrenergic receptors in pancreatic islet β cells has been extensively studied and is well-known to suppress or completely inhibit insulin secretion through Gα_i/o_-dependent signaling (Sharp 1996, Straub and Sharp 2012). While physiological circulating concentrations of catechols (epinephrine, norepinephrine) range from picomolar to low nanomolar (Clutter et al. 1980, Dodt et al. 1997, Kienbaum et al. 1998), often micromolar concentrations are used to investigate pancreatic islet function (Sieg et al. 2004, Gibson et al. 2006, Iwanir and Reuveny 2008, Zhao et al. 2008, Zhang et al. 2009, Slucca et al. 2010, Tian et al. 2011). Among the RTKs we tested in β cells, we chose TrkB for its sensitivity to stimulation with ligand, inhibition by epinephrine, and relative lack of knowledge of its role in β cells. Our characterization and analysis of BDNF-TrkB signaling to ERK1/2 in MIN6 β cells revealed effects on growth, interactions with insulin secretagogues, and that epinephrine blocks TrkB signaling at the receptor level in a G_i_-dependent manner. We conclude from our findings that the doses of epinephrine used in β cell experiments should be carefully considered when interpreting results.

## 2 Materials and Methods

### 2.1 Antibodies, Plasmids, and Reagents

All chemicals were purchased through Fisher Scientific unless otherwise indicated and listed in **Table S1**. All relevant reagents used in this study are listed in **Table S1**.

### 2.2 Plasmid generation

Mouse TrkB.FL and TrkB.T1 cDNAs were cloned from MIN6 cell cDNA libraries, generated using the Protoscript II First Strand cDNA Synthesis kit (NEB Cat# E6560S). cDNAs were inserted into pIRES-3xFlag-dsRed by restriction digestion and ligation. pX330-U6-Chimeric_BB-CBh-hSpCas9 was a gift from Feng Zhang (Addgene plasmid #42230) (Cong et al. 2013). Guide RNAs were designed targeting mouse NTRK2 exon 1 and cloned into pX330 following the Zhang lab protocol. The 20 bp guide sequences (#1: 5’-GACCCGCCATGGCGCGGCTC-3’, #2: 5’-GGAACCTAACAGCGTTGACC-3’) were generated using the CRISPR design tool (http://crispr.mit.edu). Oligonucleotides for cloning the gRNAs are shown in **Table S1**. The oligonucleotides were phosphorylated with T4 PNK and annealed for insertion into the BbsI site of pX330. All plasmids were verified by DNA sequencing (Genewiz).

### 2.3 Immunoblotting

Cleared cell lysates (40-50 μg) were separated on 10% gels by SDS-PAGE and transferred to nitrocellulose for immunoblotting. All membranes were blocked in Odyssey blocking buffer (Licor) for 1 h before overnight incubation with primary antibodies diluted in blocking buffer. After three 10 min washes in 20 mM Tris-HCl pH 7.6, 150 mM NaCl, 0.1% Tween-20 (TBS-T), membranes were incubated with fluorescent secondary antibodies for 1 h at room temperature. After three 10 min washes in TBS-T, membranes were imaged on a Licor Odyssey scanner.

### 2.4 Cell culture and transfections

MIN6 β cells were cultured in Dulbecco’s modified Eagle’s medium (D6429), supplemented with 15% fetal bovine serum, 100 units/ml penicillin, 100 μg/ml streptomycin, 292 μg/ml L-glutamine, and 50 μM β-mercaptoethanol (Kalwat et al. 2016). MIN6 cells in 12 well dishes were untreated or transfected with Lipofectamine 2000 according to the manufacturer’s instructions and cultured 48 h before use in experiments. For chronic BDNF treatment, cells were incubated with 100 ng/ml BDNF in complete culture media. Prior to stimulation, MIN6 cells were washed twice with and incubated for 2 h in freshly prepared glucose-free modified Krebs-Ringer bicarbonate buffer (MKRBB: 5 mM KCl, 120 mM NaCl, 15 mM HEPES, pH 7.4, 24 mM NaHCO3, 1 mM MgCl2, 2 mM CaCl2, and 1 mg/ml radioimmunoassay-grade BSA). For insulin secretion measurements, supernatants were collected, cleared and stored at -20°C. Cells were lysed in 25 mM HEPES, pH 7.4, 1% Nonidet P-40, 10% glycerol, 50 mM sodium fluoride, 10 mM sodium pyrophosphate, 137 mM NaCl, 1 mM sodium vanadate, 1 mM phenylmethylsulfonyl fluoride, 10 μg/ml aprotinin, 1 μg/ml pepstatin, 5 μg/ml leupeptin and cleared of insoluble material by centrifugation at 10,000 x g for 10 min at 4°C for subsequent use. Secreted and total insulin content were measured using a mouse Insulin ELISA (Mercodia).

To generate MIN6 TrkB knockout lines, 35 mm wells of MIN6 cells were co-transfected with 2.5 µg of pX330 containing gRNAs and 1 µg of pEGFP-C2 (Clontech). After 48 h, cells were trypsinized and GFP positive cells were sorted one per well in round bottom 96-well trays by fluorescence activated cell sorting for clonal expansion. After 3-4 of expansion, colonies were trypsinized and seeded into 24-well plates and screened for TrkB protein expression by immunoblotting. Clones were selected which exhibited normal morphology and low basal phosphorylated ERK1/2.

Rat INS-1 cells were cultured in RPMI-1640 containing 11 mM glucose, 10% FBS, 1 mM sodium pyruvate, 50 μM β-mercaptoethanol, 10 mM Hepes pH 7.4, penicillin/streptomycin as above. Conditions for pertussis toxin (PTX, Fisher PHZ1174) treatments have been described previously (Taussig et al. 1992, Gibson et al. 2006). Briefly, INS1 cells in 12-well culture dishes were incubated for 24 h in the presence or absence of 200 ng/ml PTX, followed by a 2 h incubation in KRBH with 2 mM glucose also in the presence or absence of 200 ng/ml PTX. Cells were then pretreated with 10 μM epinephrine for 15 min and stimulated with 10 ng/ml BDNF or 20 mM glucose for the indicated times and harvested for immunoblotting.

### 2.5 Human Pancreatic Tissue Microscopy

Paraffin-embedded formalin-fixed sections of de-identified human pancreas tissue on glass slides were obtained through the Simmons Comprehensive Cancer Center at UT Southwestern Medical. Slides were deparaffinized with the assistance of the UTSW Molecular Pathology Core using an automated system for xylene and ethanol washes. Antigen retrieval was performed by heating in citrate buffer as described (http://www.ihcworld.com/_protocols/epitope_retrieval/citrate_buffer.htm). After three 10 min washes in PBS-T (137 mM NaCl, 2.7 mM KCl, 10 mM Na_2_HPO_4_, 1.8 mM KH_2_PO_4_, pH 7.4, 0.05% Tween-20), slides were blocked for 1 h at room temperature in normal donkey serum (NDS) block solution (2% donkey serum, 1% bovine serum albumin, 0.1% cold fish skin gelatin, 0.1% Triton X-100, 0.05% sodium azide, PBS-T). Sections were outlined with a barrier pen and incubated overnight at 4°C with primary antibodies. Primary antibodies were diluted in NDS blocking solution at the indicated dilutions (**Table S1**). After three 10-min washes in PBS-T, slides were incubated in secondary antibodies in NDS block for 1 h at room temperature. The washed slides were mounted with Dapi Fluoromount-G (SouthernBiotech #0100-20) and imaged on either an LSM700 or LSM780 Zeiss confocal microscope.

### 2.6 Statistical Analysis

Quantitated data are expressed as mean ± SD. Data were evaluated using Student’s t test or ANOVA with multiple comparisons test as appropriate and considered significant if P < 0.05. Graphs were made in GraphPad Prism 8.

## 3 Results

### 3.1 Epinephrine differentially blocks activation of RTK signaling in MIN6 β cells

In our studies of β cell ERK1/2 activation, we noted an interaction between signaling downstream of RTKs and α_2_-adrenergic receptor stimulation. To expand upon these observations, we stimulated MIN6 β cells with different RTK ligands to examine the effects of epinephrine. EGF, BDNF, and FGF1 stimulated ERK1/2 phosphorylation within 5 min (**Fig 1A)**. Pretreatment with epinephrine for 15 min blocked downstream phosphorylation of ERK1/2 to varying degrees depending on the RTK in question (**Fig 1A**). We found EGF signaling to ERK1/2 was partially inhibited by epinephrine (**Fig 1A**), however BDNF- and FGF1-induced signaling appeared more sensitive. We chose BDNF-TrkB for our experiments because of its sensitivity to epinephrine and because it is relatively underexplored compared to other RTK signaling pathways in β cells.

**Figure 1.**
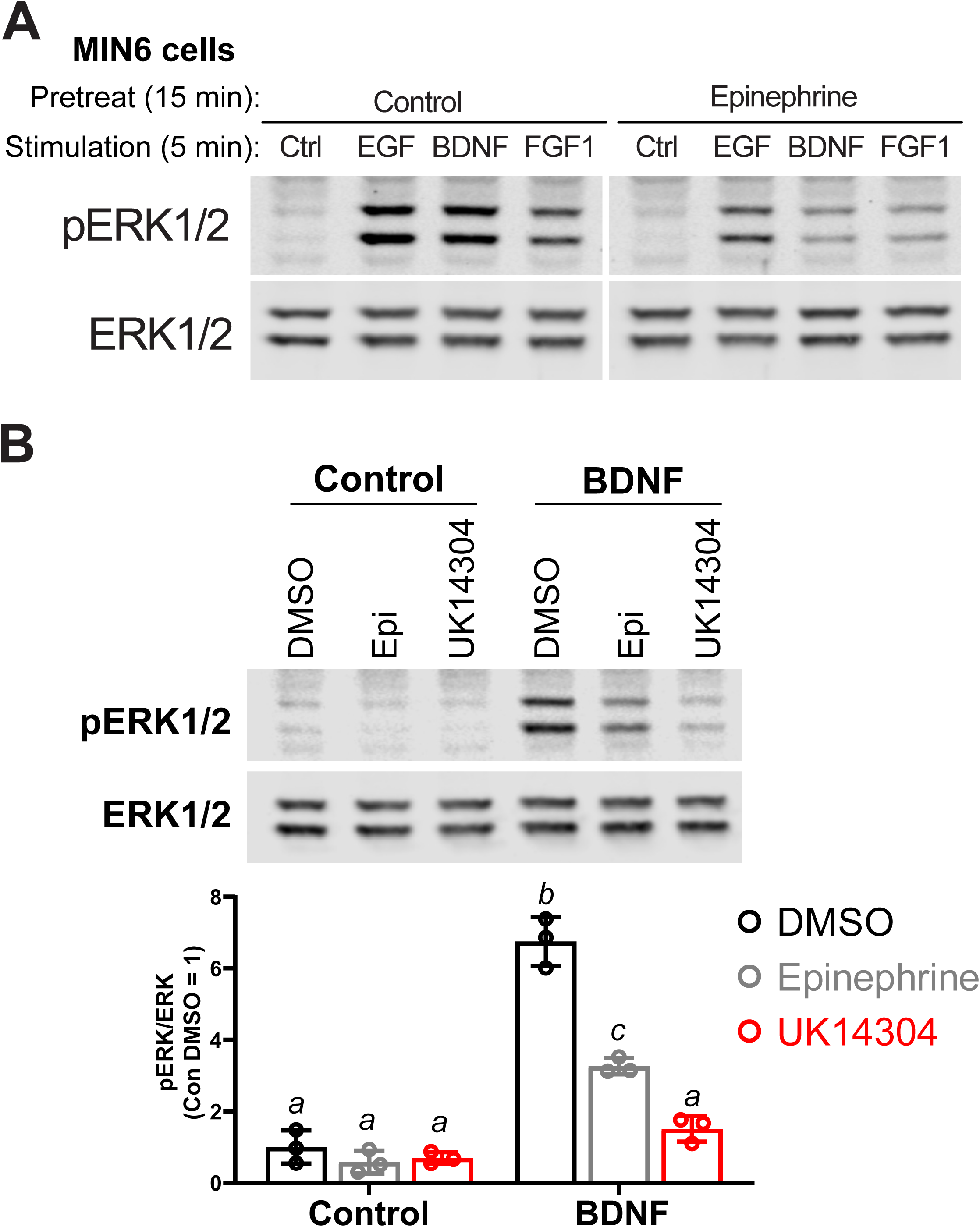
α_2_-adrenergic stimulation suppresses receptor tyrosine kinase signaling in MIN6 β cells. **A)** MIN6 cells were preincubated in KRBH for 1 h 45 min before addition of epinephrine (10 µM) for 15 min. Cells were stimulated with the indicated ligand for 5 min. (EGF 10 ng/ml; BDNF 10 ng/ml; FGF1 10 ng/ml). Immunoblots shown for phospho-ERK1/2 (pERK1/2) and total ERK1/2 are representative of three independent experiments. **B)** MIN6 cells were preincubated in KRBH for 1 h 45 min before treatment with 0.1% DMSO, 10 μM epinephrine (Epi), or 10 μM UK14304 for 15 min. Cells were then stimulated with BDNF (10 ng/ml) for 5 min. Data are the mean ± SD of three independent experiments. *, P<0.05 by 2-way ANOVA.

To confirm that epinephrine was acting through α_2_-adrenergic receptors, we tested the isoform-selective adrenergic agonist UK14304, which also suppressed BDNF-TrkB signaling to ERK1/2 in MIN6 cells (**Fig 1B**).

### 3.2 TrkB is expressed in human islets and promotes cell growth in MIN6 β cells

TrkB was reported to be expressed only in α cells (Shibayama and Koizumi 1996, Hanyu et al. 2003), however given our β cell line data we sought to confirm expression in human islets with multiple antibodies. TrkB was detected in both β and α cells in human (**Fig 2A**) and mouse (**Fig S1A**) pancreatic islets by immunocytochemistry with independently validated anti-TrkB antibodies (**Fig. S1B**). The NTRK2 gene encodes multiple isoforms of TrkB. The major forms are full-length kinase domain-containing TrkB (TrkB.FL) and a truncated form, TrkB.T1, which is missing the kinase domain (Fenner 2012). A TrkB antibody against a C-terminal epitope only found in TrkB.FL showed primarily α cell labeling (**Fig 2A**; SCBT), in agreement with previous work (Shibayama and Koizumi 1996, Hanyu et al. 2003). However, antibodies with extracellular N-terminal epitopes labeled both α and β cells (**Fig 2A**; Millipore, Abcam). We found that clonal rodent β cell lines responded to as little as 2.5 ng/ml BDNF leading to activation of ERK1/2 within 5 minutes of stimulation (**Fig. S1C-E**). BDNF had a negligible effect on the phosphorylation of Akt, but increased S6 phosphorylation at 30 min (**Fig. S1F**).

**Figure 2.**
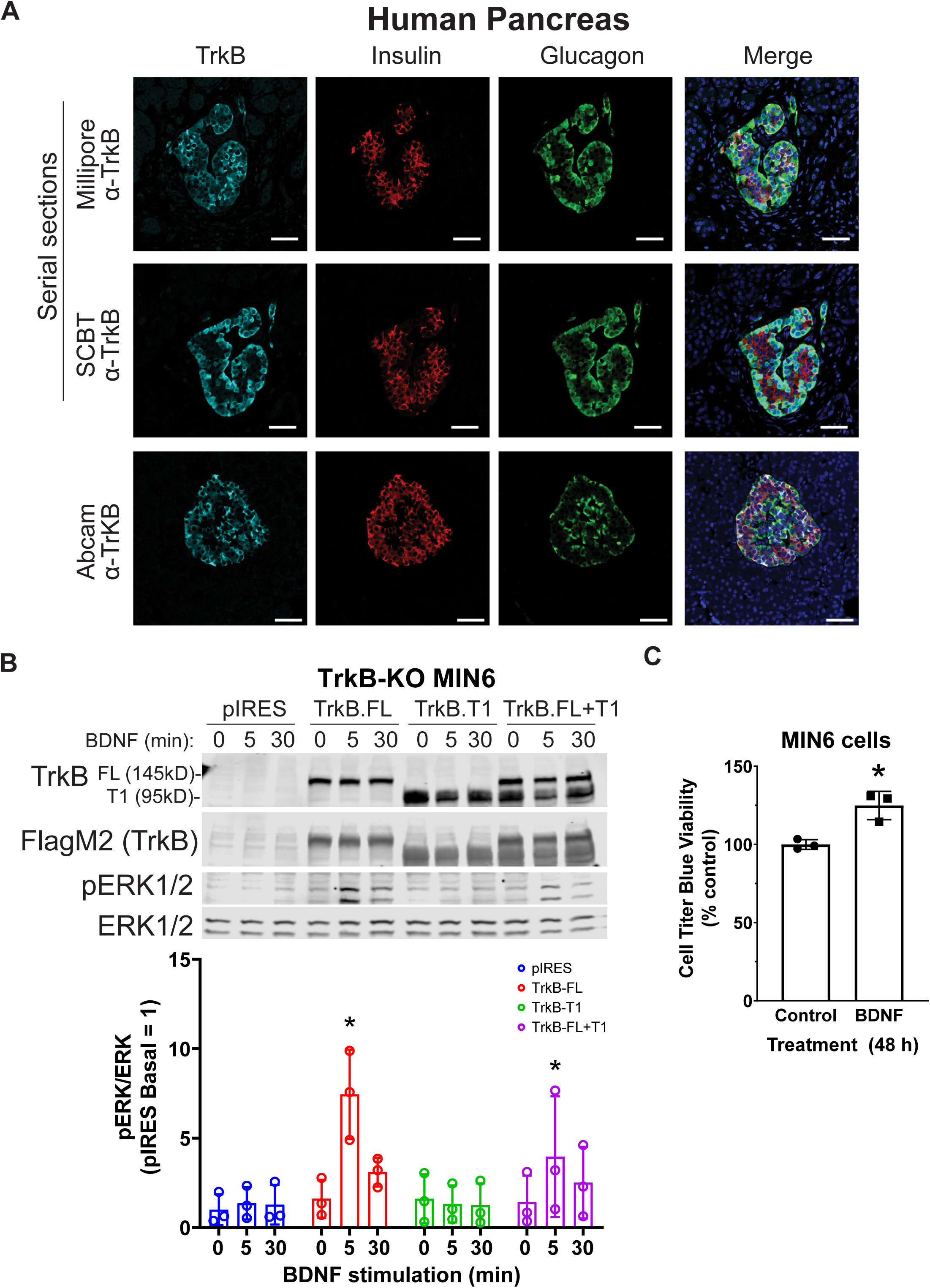
TrkB is expressed in pancreatic islets and chronic BDNF signaling promotes insulin secretion and β cell growth. **A)** Human pancreas tissue sections were immunostained with antibodies against TrkB, insulin, glucagon and also stained with DAPI. For the example shown of Millipore and Santa Cruz anti-TrkB antibody staining, serial sections from the same tissue block were stained with different TrkB antibodies and the same islet was located for imaging. Data are representative of imaging from 2 human pancreas tissue donors. Scale bar, 50 μm. **B)** TrkB KO MIN6 cells were transfected with plasmids expressing full-length TrkB (TrkB-FL), TrkB-T1, both, or empty vector. After 48 h cells were preincubated in KRBH with 4.5 mM glucose for 2 h and stimulated with 10 ng/ml BDNF for 5- and 30-min. Data are the mean ± SD of three independent experiments. *, P<0.05 for 0 vs 5 min of BDNF stimulation by two-way ANOVA using Dunnet’s multiple comparisons test. **C)** MIN6 cells plated in 96 well dishes were incubated for 48 h with BDNF (100 ng/ml) followed by Cell Titer Blue assay for viability. Bar graph is the mean ± SD from three independent passages of cells. *, P<0.05 Control vs. BDNF by t-test.

BDNF-stimulated activation of ERK1/2 was blocked by small molecule TrkB inhibitors (GNF-5837 and lestaurtinib) (**Fig. S1G-H**) as well as by CRISPR/Cas9-mediated knockout of TrkB (TrkB-KO) (**Fig. 2B**). Multiple clonal lines of TrkB knockout MIN6 cells were confirmed to lack TrkB by immunoblotting and verified to retain glucose-induced ERK1/2 activation (**Fig. S1I**). BDNF-stimulated ERK1/2 signaling was rescued upon re-expression of TrkB.FL but not TrkB.T1 (**Fig. 2B**). Additionally, we observed that 48 h of BDNF treatment increased viability (**Fig. 2C**). We did not observe any effects of chronic BDNF treatment on glucose-stimulated insulin secretion under similar conditions (**Fig. S1J**).

### 3.3 Epinephrine inhibits TrkB signaling at the receptor level and only at micromolar concentrations

To determine how α_2-_adrenergic stimulation could prevent BDNF-TrkB signaling to ERK1/2, we probed the upstream phosphorylation state of TrkB itself. Typically, BDNF stimulates autophosphorylation of the TrkB receptor at several tyrosine residues (Huang and Reichardt 2003). We found that BDNF-induced tyrosine autophosphorylation of TrkB was blocked by epinephrine in MIN6 β cells (**Fig. 3A**), raising the possibility of direct effects on the TrkB receptor tyrosine kinase. Because nanomolar concentrations of epinephrine are sufficient to inhibit glucose-stimulated insulin secretion in our InsGLuc-MIN6 reporter cells (**Fig. 3B**), we tested the ability of both 5 nM and 5 μM epinephrine to affect MIN6 responses to BDNF or EGF. 5 nM epinephrine suppressed neither BDNF or EGF signaling to ERK, nor TrkB tyrosine phosphorylation, while 5 μM epinephrine blocked BDNF signaling, yet EGF retained significant ability to activate ERK1/2 (**Fig 3C**).

**Figure 3.**
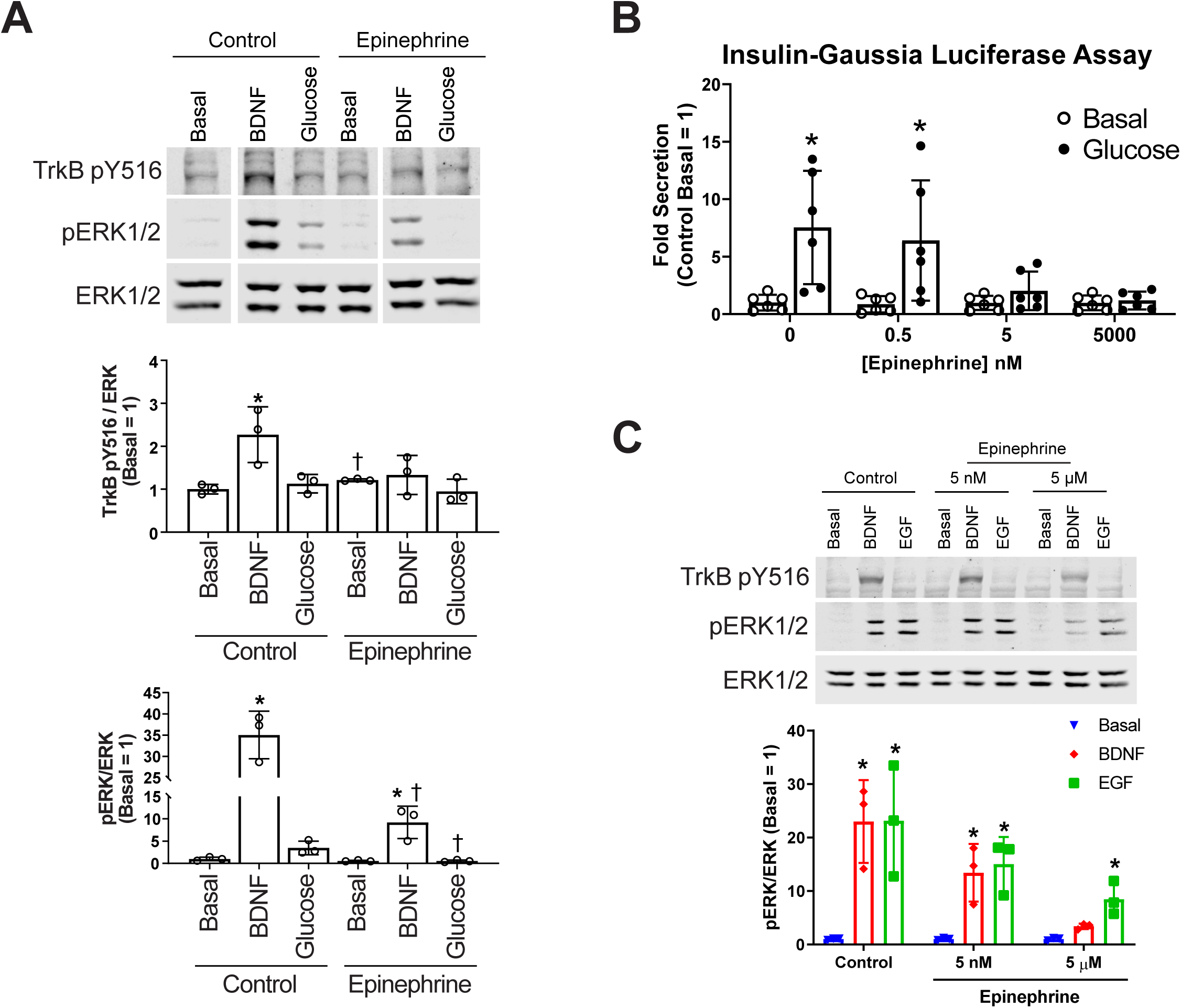
Epinephrine suppresses BDNF signaling to ERK by blocking TrkB activation. **A)** MIN6 cells were preincubated in KRBH containing 2mM glucose for 1 h 45 min before addition of epinephrine (10 µM) for 15 min. Cells were then stimulated with BDNF (10 ng/ml) or glucose (20mM) for 5 min. Western blot analysis shows phosphorylated TrkB (pY516) and ERK1/2 (pERK1/2) normalized to total ERK1/2 (N=3). *, P<0.05 vs. respective basal. †, P<0.05 Control vs. Epinephrine. **B)** MIN6 cells were treated as in (A), however the cells were pretreated with either 5 nM or 5 µM epinephrine prior to stimulation with EGF or BDNF for 5 min. Western blot analysis shows only 5 µM epinephrine suppressed pTrkB and pERK1/2 signaling by BDNF (N=3). *, P<0.05. **C)** InsGLuc-MIN6 cells were preincubated in glucose-free KRBH for 1 h followed by stimulation with glucose (20 mM) in the presence or absence of epinephrine (0.5 nM, 5 nM, or 5 µM) for 1 h. Supernatant was collected for Gaussia luciferase assays and the data are reported as the fold change with respect to basal unstimulated cells (N=6). *, P<0.05 Basal vs. Glucose. All graphs are the mean ± SD.

### 3.4 Epinephrine-mediated inhibition of BDNF-TrkB signaling depends on G_i_ but does not involve calcium influx or cAMP generation

Epinephrine inhibits adenylyl cyclase and insulin secretion in β cells through α_2_-adrenergic receptors linked to Gα_i_ (Straub and Sharp 2012). To delve further into the mechanism of epinephrine-mediated blockade of BDNF-TrkB signaling, we treated cells with pertussis toxin (PTX) which inactivates Gα_i_. INS1 β cells were used because in our experience they exhibit a more robust response to PTX than MIN6 cells. PTX rescued the effects of epinephrine on BDNF- and glucose-mediated activation of ERK1/2 (**Fig. 4A**), indicating a requirement for Gα_i/o_ signaling. We also examined the involvement of calcium influx, which regulates ERK1/2 activation in β cells and is inhibited by epinephrine (Straub and Sharp 2012). Eliminating calcium influx by removal of calcium from the incubation buffer had no impact on BDNF signaling to ERK1/2, nor did it prevent the inhibiting effect of epinephrine on BDNF-induced ERK1/2 activation. KCl-mediated depolarization no longer activated ERK1/2 in the absence of calcium (**Fig. 4B**), demonstrating the mechanistic dichotomy between growth factor and depolarization-stimulated ERK1/2 activation.

**Figure 4.**
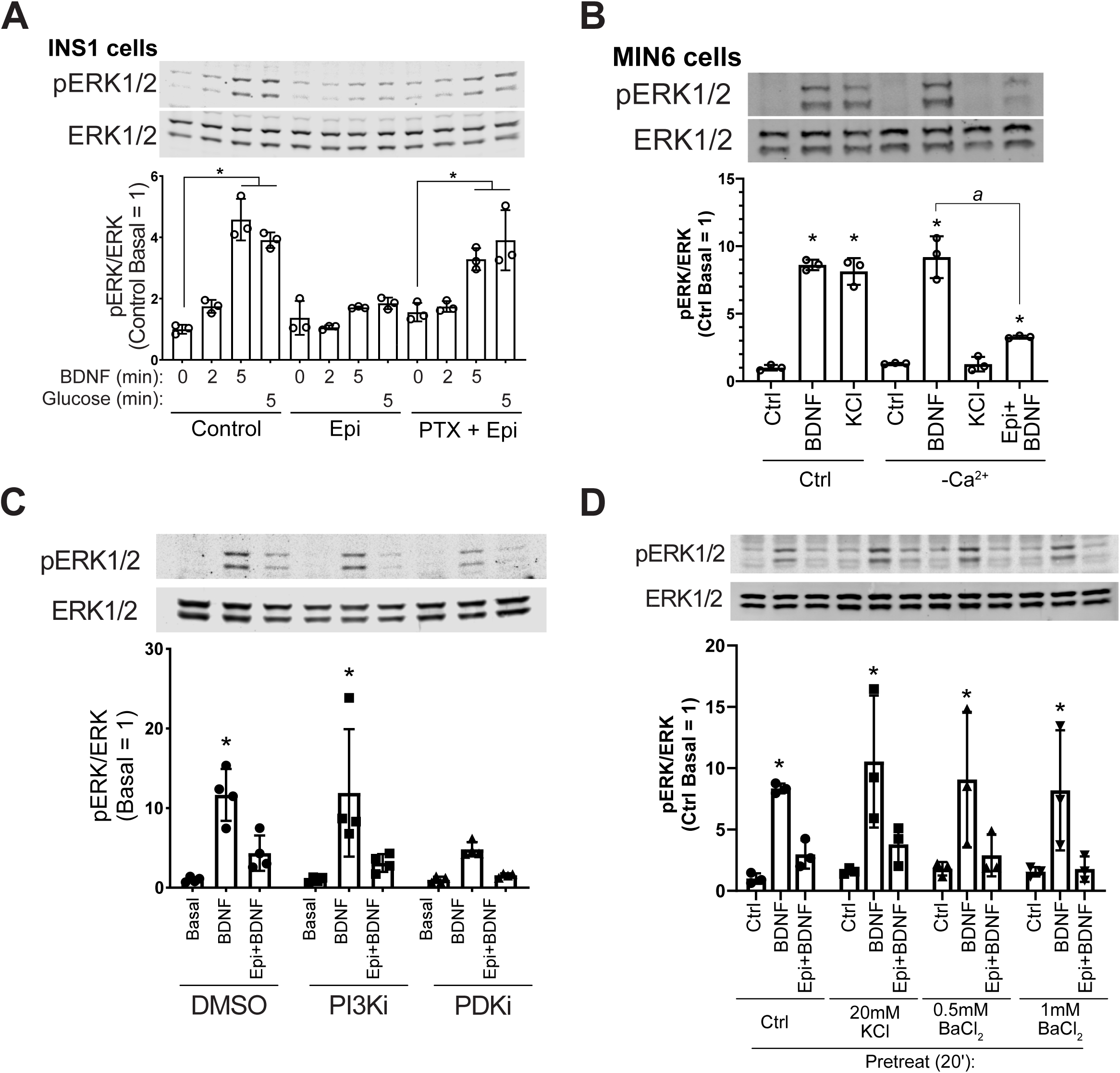
Epinephrine blockade of BDNF signaling depends on Gα_i/o_, but is unaffected by Ca^2+^ influx, PI3K/PDK1 inhibition, or treatment with KCl or BaCl_2_. **A)** INS1 β cells were treated with 200 ng/ml pertussis toxin (PTX) for 18 h in culture medium. Cells were then incubated in KRBH with 2 mM glucose in the continued presence or absence of PTX for 2 h. Prior to stimulation, cells were treated with or without 10 µM epinephrine for 15 min. Cells were stimulated with or without 10 ng/ml BDNF or 20 mM glucose for the indicated time. Data are the mean ± SD of three independent experiments. *, P<0.05 vs respective Basal. **B)** MIN6 cells were preincubated in normal (Ca^2+^-containing) or Ca^2+^-free KRBH (compensated with additional 2 mM MgCl_2_) without glucose for 2 h. 15 min prior to stimulation 10 μM epinephrine was added where indicated. Cells were stimulated with 10 ng/ml BDNF or 50 mM KCl for 5 min. Data are the mean ± SD. *, P<0.05 compared to respective basal. *a*, P<0.05. **C)** MIN6 cells were preincubated for 1.5 h in glucose-free KRBH and then treated with DMSO (0.1%), GDC-0941 (250 nM) or GSK2334470 (250 nM) for 15 min. Cells were then treated or not with 5 μM epinephrine for 15 min before stimulation with BDNF (10 ng/ml) for 5 min. Bar graph represents the mean ± SE for four independent experiments. *, P<0.05 vs. Basal by two-way ANOVA with Dunnett’s multiple comparisons test. **D)** MIN6 cells were preincubated in glucose-free KRBH for 1 h and 40 min at which point KCl or BaCl_2_ were added. At 1 h and 45 min, epinephrine (5 µM) was added. At 2 h cells were stimulated with BDNF for 5 min and then harvested for Western blot analysis. All data are the mean ± SD of N=3 experiments. *, P<0.05.

Circulating BDNF concentrations are altered in diabetes and BDNF acts through multiple pathways present in β cells. Cross-talk between receptor tyrosine kinases and GPCRs has been found (Marty and Ye 2010), although little is known about these pathway interactions in β cells. We tested whether BDNF exhibits crosstalk not only with epinephrine, but also with glucagon-like peptide 1 (GLP-1) and glucose. Pre-treatment with GLP-1 enhanced the ERK1/2 response to the combination of BDNF and glucose (**Fig. S2A**), suggesting interactions among TrkB, the GLP-1 receptor, and glucose metabolic pathways. Epinephrine pretreatment dramatically inhibited ERK1/2 activation in response to either glucose, BDNF or their combination. Because of the potential for interaction with the GLP1 receptor, we tested whether BDNF can induce cAMP generation in MIN6 cells expressing a bioluminescence-resonance energy transfer-based cAMP reporter (<u>cAM</u>P sensor using <u>Y</u>FP-<u>E</u>pac-RL<u>u</u>c or CAMYEL) (Jiang et al. 2007, Guerra et al. 2017). While the known G_s_ activator GLP-1 increased cAMP (Holst 2007), BDNF had no effect on cAMP, either alone or in combination with glucose or GLP-1 (**Fig. S2B**).

### 3.5 Epinephrine inhibition of RTKs is not prevented by PI3K/PDK1 inhibition or by blockade of potassium channels with KCl or BaCl_2_

Inhibition of the phosphoinositide-3-kinase signaling pathway is reported to abrogate the hyperpolarizing effects of micromolar epinephrine (Zhang et al. 2009). We pretreated MIN6 β cells with well-characterized inhibitors for PI3K (GDC-0941) and PDK1 (GSK2334470) followed by epinephrine and analyzed the response to BDNF. These inhibitors did not rescue BDNF-induced ERK1/2 activation (**Fig 4C**), suggesting the PI3K/PDK1 pathway is not required for the G_i_-dependent blockade of TrkB activation.

Sieg et al. found that epinephrine hyperpolarizes cells in a PTX-sensitive manner through unidentified K^+^ channels that can be blocked by low dose (20 mM) KCl or 0.5-1 mM BaCl_2_ (Sieg et al. 2004). We tested these conditions and found that treating with KCl or BaCl_2_ prior to addition of micromolar epinephrine did not rescue BDNF-TrkB signaling to ERK1/2 (**Fig 4D**). Another possible explanation is that epinephrine somehow induces TrkB receptor internalization. However, surface biotinylation experiments did not show a significant effect of epinephrine on the amount of surface TrkB (**Fig S3**), suggesting alternative mechanisms.

## 4 Discussion

### 4.1 How does α_2_-adrenergic stimulation block activation of RTKs like TrkB?

In addition to the events that distinguish signaling by BDNF and glucose, the unexpected sensitivity of BDNF to inhibition by α_2_-adrenergic agonism also suggests a connection between signaling by BDNF and insulin secretagogues. In general, RTKs share certain signaling pathways; the Ras-ERK1/2, PI3K-Akt and PLCγ pathways are the most recognized (Minichiello 2009). Depending on context, different ligand-receptor family members may signal independently within the same cell to different pathways or exhibit inter-pathway crosstalk (Coster et al. 2017), but the exact molecular mechanisms are not always clear. Additionally, other inputs such as glucose-stimulated metabolic pathways can act on some of the same signaling pathways seemingly by independent mechanisms. (Khoo and Cobb 1997, Khoo et al. 2004, Kalwat et al. 2013). α_2_-adrenergic signaling is well-known to antagonize insulin secretion in β cells, largely through heterotrimeric G_i/o_ proteins (Gibson et al. 2006, Zhao et al. 2010, Straub and Sharp 2012, Ito et al. 2017). In tandem with this effect, α_2_-adrenergic signaling blocks glucose stimulated ERK1/2 phosphorylation (Gibson et al. 2006), by mechanisms presumably including membrane hyperpolarization, inhibition of adenylyl cyclase and blockade of calcium influx. We found that BDNF signaling through the Raf-MEK-ERK pathway is also blocked in β cells by epinephrine through a G_i_-dependent mechanism at the level of TrkB activation, indicating that essential β cell regulatory inputs are shared between RTKs and glucose-stimulated signaling pathways.

G_i_-dependent crosstalk between α_2_-adrenergic receptors and RTKs is a relatively unexplored aspect of β cell signaling. GPCR-RTK crosstalk has been observed to activate RTK pathways, but reports of G protein inhibition of RTKs are uncommon (Marty and Ye 2010). Endogenous plasma epinephrine concentrations in humans are usually 10-100 pg/ml (54.5-545 pM) (Dodt et al. 1997, Kienbaum et al. 1998), but can increase to 1000 pg/ml (5.4 nM) (Clutter et al. 1980) during infusions of drugs or epinephrine itself. High (μM) and low (nM) doses of epinephrine have been proposed to inhibit insulin secretion through different mechanisms (Ito et al. 2017). In the case of nanomolar concentrations of epinephrine, cAMP-TRPM2 channel activity is suppressed, blunting glucose-induced insulin secretion, although sulfonylurea-induced secretion is unaffected (Ito et al. 2017). At ≥1 μM epinephrine, secretion under nearly all conditions is inhibited and the plasma membrane is hyperpolarized. In work from Peterhoff, et al. (Peterhoff et al. 2003), 1 μM epinephrine was shown to inhibit adenylyl cyclase and hyperpolarize the plasma membrane in wild-type β cells, but not in α_2A_-adrenergic receptor knockout β cells, suggesting the actions of micromolar concentrations of epinephrine occur specifically through its receptor.

Because activation of RTK signaling by BDNF in MIN6 β cells appears unaffected by changes in intracellular calcium and does not induce cAMP on its own, suppression of these mechanisms is unlikely to account for the effect of epinephrine on BDNF-TrkB signaling. However, we found that micromolar concentrations of epinephrine were required for its inhibitory activity on TrkB. Why nanomolar epinephrine fails to block BDNF-TrkB signaling when it is also known to activate G_i/o_ under those conditions is an open question. Possible mechanisms may include membrane hyperpolarization or activation of G protein-gated inward rectifier potassium channels (Iwanir and Reuveny 2008), although it is unclear how membrane polarization could affect TrkB. One possibility is that G_i/o_ binds directly to TrkB to inhibit its activation as was shown for G_i/o_ binding to insulin receptor in β cells (Kim et al. 2012); however, mechanisms for concentration-dependent effects of epinephrine in such a process are unclear.

### 4.2 What is the role for BDNF-TrkB signaling in β cells?

BDNF and its receptor TrkB mediate aspects of neuronal development and differentiation and are essential in whole body energy homeostasis (Hutchison 2012, Hutchison 2013) and are implicated in diabetes (Verge et al. 2014). BDNF can indirectly regulate islet hormones, like glucagon, through actions in the hypothalamus and innervation of the islet (Gotoh et al. 2013) and was suggested to have direct effects on α cells (Hanyu et al. 2003). However, TrkB is expressed in both rodent and human β and α cells, as supported by our data and others (Shibayama and Koizumi 1996, Hanyu et al. 2003, Uhlen et al. 2015, DiGruccio et al. 2016, Segerstolpe et al. 2016, Fulgenzi et al. 2020). TrkB is expressed as two splice isoforms, full-length TrkB and TrkB.T1 which is missing the cytosolic kinase domain, instead containing a short distinct cytosolic tail (Fenner 2012). Recent work from Fulgenzi, et al. has demonstrated that the splice isoform of TrkB.T1 is involved in BDNF-induced insulin secretion (Fulgenzi et al. 2020). While TrkB.T1 expression is much greater than full-length TrkB in β cells, full-length TrkB is expressed and even a relatively low amount of RTK at the protein level is sufficient for signaling. We have observed that even >90% knockdown of TrkB protein by siRNA was insufficient to blunt BDNF stimulated ERK1/2 activation. Not until TrkB protein was eliminated completely by CRISPR/Cas9 did we prevent BDNF-ERK1/2 signaling.

There are multiple studies linking circulating BDNF concentration to type 2 diabetes in humans and mice (Krabbe et al. 2007, Sha et al. 2007, Li et al. 2016, Murillo Ortiz et al. 2016) as well as in type 1 diabetic patients (Tonoli et al. 2015). BDNF null animals have elevated blood glucose and usually die by the second post-natal week (Hanyu et al. 2003). Treating db/db mice with BDNF lowered blood glucose and increased pancreatic insulin content (Tonra et al. 1999). β cell area and staining intensity were increased and pancreatic glucagon content was decreased (Yamanaka et al. 2006). BDNF may also have a cytoprotective role in the islet because treatment with BDNF prevented RIN5F β cell death in response to alloxan, streptozotocin, doxorubicin and benzo(a)pyrene (Bathina et al. 2016). Additionally, we observed that the stimulatory concentration of BDNF is well within the range of the circulating hormone (Krabbe et al. 2007, Dell’Osso et al. 2009, Matthews et al. 2009, Karczewska-Kupczewska et al. 2012, Kurita et al. 2012, Pillai et al. 2012). These studies indicate the need for further analysis of the effects and mechanisms of action of BDNF-TrkB signaling in pancreatic islets.

TrkB is known to exhibit crosstalk with other kinases, including Src-family kinases (Huang and McNamara 2010), Ret (Esposito et al. 2008) and the EGF receptor (Puehringer et al. 2013). Oligomerization between receptor kinases TrkA and TrkB could potentially be a contributor to the actions of NGF and may also contribute to BDNF-TrkB signaling not reflected by ERK1/2 activity. Another factor that may complicate interpretation of BDNF function is the expression of isoforms lacking the kinase domain. In our knockout-rescue experiments TrkB.T1 seemed to suppress the activity of the full length receptor, as has been suggested in other systems (Eide et al. 1996, Fryer et al. 1997, De Wit et al. 2006). In the future. specific deletion of TrkB.FL from different islet cell types in mice from early in development or in the adult may deconvolute roles for TrkB.FL and TrkB.T1 in islet development and function.

### 4.3 Future directions

Whether epinephrine or G_i/o_-activation impacts RTK signaling in other cell types is an open question. Adrenergic stimulation of β cells has been suggested to impair β cell growth at near micromolar concentrations (Zhao et al. 2014), and while cAMP is potentially involved, other mechanisms including suppression of RTK signaling may be at work in such conditions.

Future studies are required to place these actions of BDNF in an *in vivo* context to assess their metabolic impact, as well as to elucidate the mechanism underlying the unanticipated finding that epinephrine prevents activation of TrkB itself by BDNF. If any components of that mechanism are pharmacologically targetable, it may be possible to modulate TrkB or other RTK signaling in islets *in vivo* for therapeutic benefit. Notably, single nucleotide polymorphisms in and near the ADRA2A gene (encoding α_2A_-adrenergic receptor) have been correlated with increased fasting glycemia and type 2 diabetes risk (Liggett 2009, Dupuis et al. 2010, Talmud et al. 2011, Langberg et al. 2013); impaired glucose-stimulated insulin secretion is a factor in this increased risk. Our findings suggest there is potential for contribution of altered adrenergic crosstalk with RTK signaling in disease. In addition, these results have implications for other systems in which adrenergic and receptor tyrosine kinase signaling may converge, such as cancer (Hui et al. 2008, Powe et al. 2011, Stock et al. 2013).

## 7 Author Contributions

Conceptualization, MAK; Formal Analysis, MAK; Investigation, MAK, ZH, DB, KM; Writing-Original draft, MAK, MHC; Supervision, MAK, MHC; Funding Acquisition, MAK, MHC.

## 8 Funding

NIH F32 DK100113 and JDRF 2-SRA-2019-702-Q-R to MAK.

NIH R01 DK55310, NIH R37 DK34128, and Welch I1243 to MHC.

UTSW Simmons Comprehensive Cancer Center, NCI P30 CA142543, supports multiple core services.

## 9 Acknowledgments

Thank you to members of the Cobb and Albanesi labs for advice. Specifically thank you to Magdalena Grzemska and Ji-ung Jung for critical reading of this manuscript and to Dionne Ware for administrative assistance. We thank the lab of James Collins for use of microscopes. We thank the UTSW Simmons Comprehensive Cancer Center (NCI P30 CA142543) cores for Live Cell Imaging, Tissue Management Shared Resource, and the UTSW histopathology core. Thanks to the UTSW Flow Cytometry facility. Thanks to the lab of Louis Parada for the 3T3-TrkB cell line. During the course of this work, MAK was supported by an NIH NRSA DK100113 and a JDRF 2-SRA-2019-702-Q-R. ZH was supported by the UT Southwestern-Sun Yat-Sen exchange program. Also supporting this work were R01 DK55310, R37 DK34128 and grant I1243 from the Welch Foundation to MHC. This manuscript has been released as a pre-print at bioRxiv.

## 11 Supplementary Material

Supplementary figures, table, and legends are provided separately and available online.

