## Supplemental Materials for "α_2_-adrenergic signaling disrupts β cell BDNF-TrkB receptor tyrosine kinase signaling"

### *Supplementary Material*

#### 1 Supplementary Methods

##### 1.1 Microscopy of mouse pancreas and NIH-3T3-TrkB cells

Paraffin-embedded pancreas tissue sections from db/db mice were probed for TrkB expression using the Millipore antibody, as with human pancreas tissue sections. We tested TrkB antibodies using NIH-3T3-TrkB cells (Soppet et al. 1991) by immunofluorescence microscopy. We determined that antibodies from Millipore and Santa Cruz Biotechnologies detected human TrkB. Two other antibodies from Santa Cruz Biotechnology and Origene were not optimal. In other immunoblotting experiments we observed that rabbit anti-TrkB from Abcam (Cat# ab134155, 1:1000) also detected TrkB.FL in differentiated SHSY5Y cells.

##### 1.2 Intracellular cAMP measurements

We used the Epac-based bioluminescence resonance energy transfer (BRET) sensor for cAMP (CAMYEL) in MIN6 cells as previously described (Jiang et al. 2007, Guerra et al. 2017). Briefly, BRET assays were performed on the Synergy H1 microplate reader (BioTek). MIN6 CAMYEL cells were preincubated in KRBH without glucose for 1.5 h followed by addition of 2.55 μM coelenterazine-h for 15 min. Assays were performed in triplicate. Emission signals at 485/20 and 528/20 nm (center/bandpass) were measured every 0.8 s during a 3 min baseline measurement. Stimuli were then added and cells were measured for an additional 10 min.

##### 1.3 Surface Biotinylation Assay

Sulfo-NHS-SS-biotin was resuspended in DMSO and stored in single-use 400X (200 mg/mL) aliquots at -20°C. MIN6 cells were preincubated in glucose-free KRBH without BSA for 1.75 h prior to treatment with or without 10 μM epinephrine for 15 min. KRBH was removed, cells placed cold shaker in 4°C walk-in fridge and cold PBS was added containing with Sulfo-NHS-SS-biotin (0.5 mg/ml) for 30 min to label surface exposed proteins. 3 x 10 min washes with 0.1M glycine to stop the reaction. Cells were lysed in lysis buffer (2% NP-40, 150 mM NaCl, 0.01% SDS, 50 mM HEPES pH 7.5 and protease/phosphatase inhibitors) and 2 mg cleared lysates were used in streptavidin-agarose pulldowns. The precipitated proteins were run on SDS-PAGE and immunoblotted for TrkB. ATP1A1 and ADRA2A were also blotted as controls. Surface TrkB and ADRA2A were normalized to surface ATP1A1.

#### 2 Supplementary Figures

##### 2.1 Supplementary Figures

**Supplementary Figure 1. TrkB is expressed in pancreatic islets and BDNF stimulates the canonical TrkB RTK signaling pathway in β cells.**

**A)** db/db mouse pancreas tissue sections were immunostained for TrkB, insulin, and glucagon. Scale bar, 20 μm.

**B)** Anti-TrkB immunofluorescent staining in 3T3-TrkB cells to validate antibody suitability.

**C)** INS1 cells were preincubated 2 h in glucose-free KRBH and then stimulated with glucose (20 mM) or BDNF (10 ng/ml) for 5 min.

**D,E)** MIN6 cells were preincubated in glucose-free KRBH and then treated with increasing doses of BDNF for 5 min. pERK1/2 and ERK1/2 were immunoblotted and quantitation is show below as bar graphs. Data are the mean ± SE of three independent passages of cells. *, $P<0.05$.

**F)** MIN6 cells were preincubated in KRBH with 2 mM glucose for 2h and then treated with or without 10 ng/ml BDNF for 5 and 30 min. Data are the mean ± SE of three independent experiments. *, $P<0.05$.

**G)** Time course of BDNF treatment in MIN6 cells at 10 and 100 ng/ml. High dose BDNF leads to TrkB-FL degradation.

**H,I)** MIN6 cells were preincubated in KRBH with 2 mM glucose for 2 h prior to a 10 min pretreatment with either GNF-5837 (10 µM) or lestaurtinib (Lest, 2 µM) prior to stimulation with BDNF (50 ng/ml).

**J)** One wild type and two distinct clonal lines of TrkB KO (KO1, KO2) MIN6 cells were preincubated in KRBH without glucose for 2 h and stimulated with 20 mM glucose for 5 min. Data are the mean ± SE of three experiments.

**K)** MIN6 cells were incubated with BDNF (100 ng/ml) for 24 h followed by incubation in KRBH to assay for glucose-stimulated insulin secretion. Insulin secretion (as % content) was normalized to basal secretion. Bar graph is the mean ± SD from three independent passages of cells. *, $P<0.05$ for 0 vs 30 min by two-way ANOVA using Dunnet's multiple comparisons test.

**Supplementary Figure 2. BDNF-TrkB exhibits crosstalk with GLP-1 and adrenergic signaling pathways in the β cell, but does not induce cAMP.**

**A).** MIN6 cells were preincubated in glucose-free KRBH for 2 h with or without the addition of 10 ng/ml BDNF (150 min final treatment). Prior to the glucose addition cells were treated with or without 50 nM GLP-1 or 5 µM epinephrine for 15 min. After 2 h, 20 mM glucose was added either alone, to the BDNF pre-treated cells, or in combination with BDNF for 30 min. Phosphorylated ERK1/2 (pERK1/2) and total ERK1/2 immunoblots are shown and the quantitated ratio of pERK/ERK is shown below. Bar graph is the mean ± SD of three independent experiments.

**B)** Intracellular cAMP generation was measured using MIN6 CAMYEL cells. Cells were preincubated in KRBH without glucose for 1.5 h and treated with 20 µM coelenterazine for 20 min. After a 3 min baseline measurement, stimuli were injected and BRET signal was monitored. Stimuli concentrations were: 20 mM Glucose, 30 nM GLP-1, 10 ng/ml BDNF. Data are the mean ± SE of three experiments performed in triplicate.

**Supplementary Figure 3. TrkB internalization was unaffected by epinephrine.**

MIN6 cells were preincubated in KRBH without BSA for 2 h and then stimulated with epinephrine (10 µM) for 15 min. Immediately after, cells were transferred to ice and labeled with NHS-SS-biotin to biotinylate surface proteins, the reaction was quenched with glycine and lysates were prepared for streptavidin bead precipitation and Western blot analysis. The $\alpha_2$-adrenergic receptor and TrkB bands were normalized to Na/K-ATPase (ATP1A1) and to the basal unstimulated condition. Bar graph is mean ± SE for N=3-4.

#### 4 Supplementary Table

**Supplementary Table 1**. All antibody information including RRIDs are provided. Antibodies for immunoblotting were diluted in Licor Blocking buffer (diluted 1:5 in TBS-T). Primary incubation was overnight. Secondary incubation was for 1h. Blots were washed 3 times for 10min each after primary and secondary. For immunofluorescence all blocking was 1 h at RT, primary overnight at 4°C and secondary 1h RT. All washing was 3 times for 10min each in PBS-T at RT. IB, immunoblotting; IF, immunofluorescence. All specialized reagents, primers, and tissue donor information is also provided.

**Antibody Details**

| Antigen | Host | Mol Weight (kDa) | Company | Cat# | RRID | Dilution for IB | Dilution for IF | Epitope |
|---|---|---|---|---|---|---|---|---|
| pERK1/2 | Mouse | 42,44 | Cell Signaling Technologies | 9106 | AB_331768 | 1:1000 | - | |
| ERK1/2 | Rabbit | 42,44 | In-house | Y691 | | 1:3000 | - | |
| TrkB | Rabbit | 145, 95 | Millipore | 07-225 | AB_310445 | 1:1000 | 1:100 | extracellular domain of rat TrkB |
| TrkB | Rabbit | 145, 95 | Abcam | ab134155 | AB_2857962 | - | 1:100 | Residues 130-160 in human TrkB |
| TrkB | Rabbit | 145, 95 | Santa Cruz Biotechnologies | sc-12 | AB_632557 | - | 1:100 | Residues 760-810 of mouse TrkB |
| TrkB | Mouse | 145, 95 | Origene | TA500386 | AB_2155134 | - | 1:100 | |
| TrkB | Mouse | 145, 95 | Santa Cruz Biotechnologies | sc-377218 | AB_2801499 | | 1:100 | |
| pY490/516 TrkA/B | Rabbit | 145 | Cell Signaling Technologies | 4619 | AB_10235585 | 1:1000 | - | |
| Insulin | Goat | 6 | Santa Cruz Biotechnologies | sc-7839 | AB_2296108 | - | 1:300 | |
| Glucagon | Mouse | 3.4 | Sigma | G2654 | AB_259852 | - | 1:1000 | |
| Akt | Mouse | 60 | Cell Signaling Technologies | 2967 | AB_331160 | 1:1000 | - | |
| pS473 Akt | Rabbit | 60 | Cell Signaling Technologies | 4060 | AB_2315049 | 1:1000 | - | |
| S6 (pS235/244) | Rabbit | 32 | Cell Signaling Technologies | 2211 | AB_331679 | 1:1000 | - | |
| S6 | Mouse | 32 | Santa Cruz Biotechnologies | sc-74459 | AB_1129205 | 1:3000 | - | |
| ATP1A1 | Mouse | 112 | Invitrogen | MA1-16731 | AB_2060993 | 1:1000 | - | |
| ADRA2A | Rabbit | 51 | Proteintech | 14266-1-AP | AB_2636822 | 1:1000 | - | |

**Secondary Antibodies (IB)**

| | | | | | | | |
|---|---|---|---|---|---|---|---|
| anti-rabbit 680 (red) | Donkey | | Licor | 92668073 | AB_10954442 | 1:10,000 | - |
| anti-mouse 800 (green) | Donkey | | Licor | 92632212 | AB_621847 | 1:10,000 | - |

**Secondary Antibodies (IF)**

| | | | | | | | |
|---|---|---|---|---|---|---|---|
| anti-rabbit 647 | Donkey | | Invitrogen | A-31573 | AB_2536183 | - | 1:400 |
| anti-goat 555 | Donkey | | Invitrogen | A-21432 | AB_141788 | - | 1:400 |
| anti-mouse 488 | Donkey | | Invitrogen | A-21202 | AB_141607 | - | 1:400 |

**Reagents and other materials**

| **Name** | **Company** | **Catalog #** |
|---|---|---|
| GLP-1 | Sigma | G8147 |
| Lipofectamine 2000 | Fisher | 11668019 |
| UK14304 | Sigma | U104 |
| EGF | Gibco | PHG0311 |
| FGF1a | Biolegend | 750902 |
| BDNF | Millipore | GF029 |
| Pertussis Toxin (PTX) | Fisher | PHZ1174 |
| Epinephrine | Sigma | E4375 |
| GSK2334470 | Apexbio | B2174 |
| GDC-0941 | Apexbio | A8210 |
| $BaCl_2$ | Sigma | 202738 |
| Cell Titer Blue | Promega | G8080 |
| Round-bottom 96 well culture dishes | Corning | 3799 |
| Mouse Insulin ELISA | Mercodia | 10-1247-10 |
| Sulfo-NHS-SS0-biotin | Fisher | PI21331 |

| | | |
|---|---|---|
| coelenterazine-h | Fisher | PR-S2011 |

**Primers**

| Name | Sequence (5' -> 3') |
|---|---|
| NTRK2-#1_sense | CACCGACCCGCCATGGCGCGGCTC |
| NTRK2-#1_antisense | AAACGAGCCGCGCCATGGCGGGTC |
| NTRK2-#2_sense | CACCGGAACCTAACAGCGTTGACC |
| NTRK2-#2_antisense | AAACGGTCAACGCTGTTAGGTTCC |

**Human Pancreatic Donor Tissue Information**

| UTSW ID# | Diagnosis |
|---|---|
| 18037 | Mucinous Carcinoma of the panc (tail) |
| 3394 | Intraductal papillary mucinous neoplasm |
