## Supplementary figures and images for "α_2_-adrenergic signaling disrupts β cell BDNF-TrkB receptor tyrosine kinase signaling"

### Figure S1

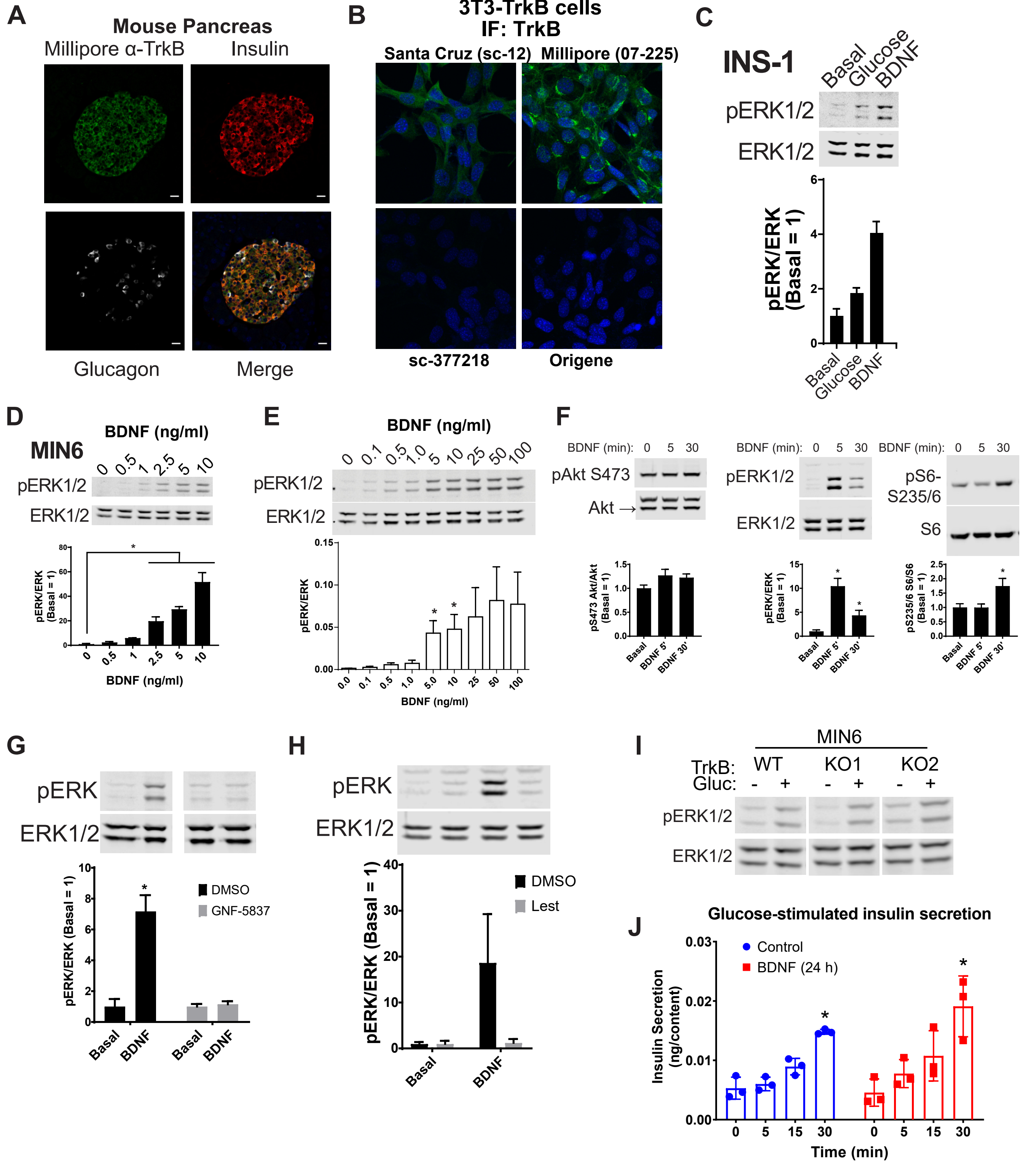

### Figure S2

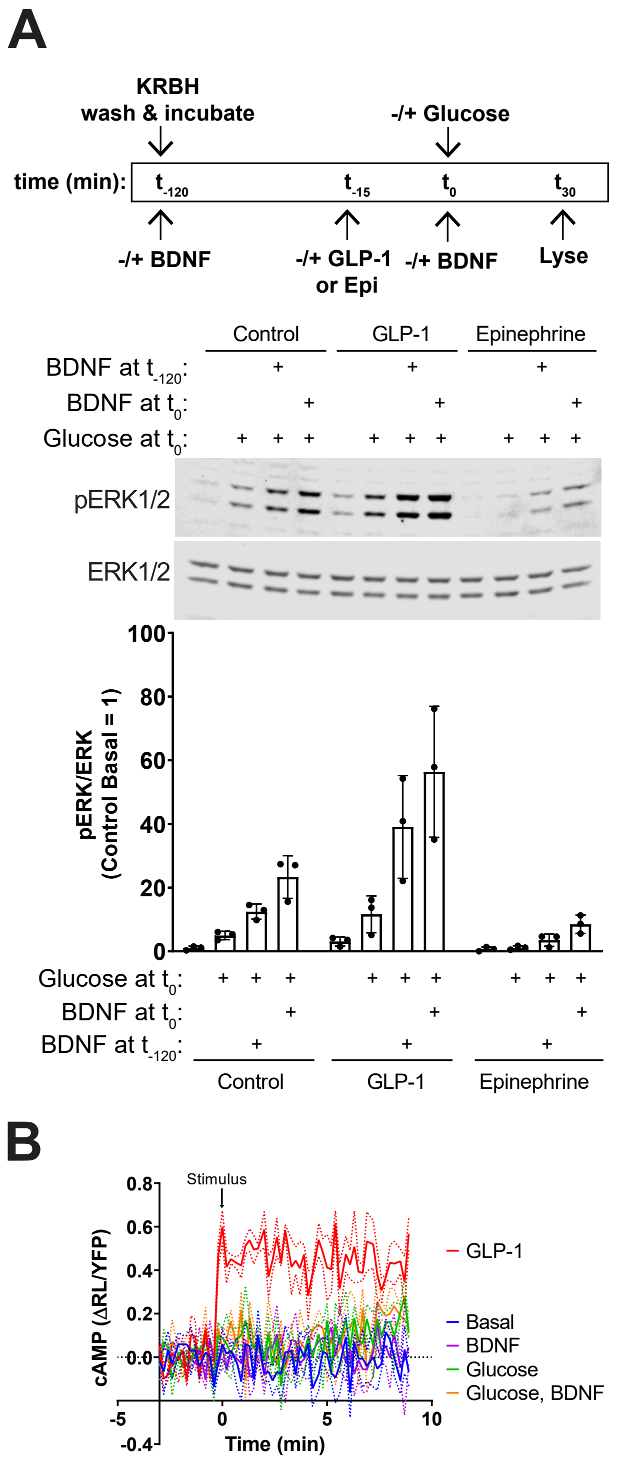

### Figure S3

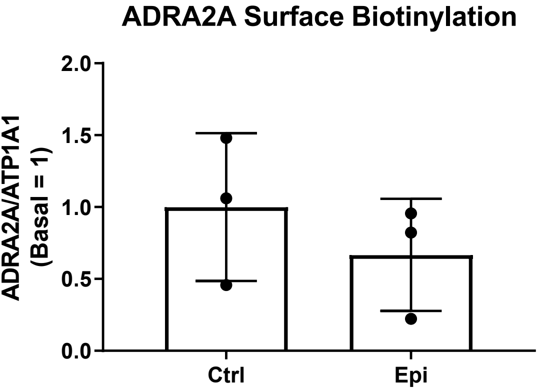

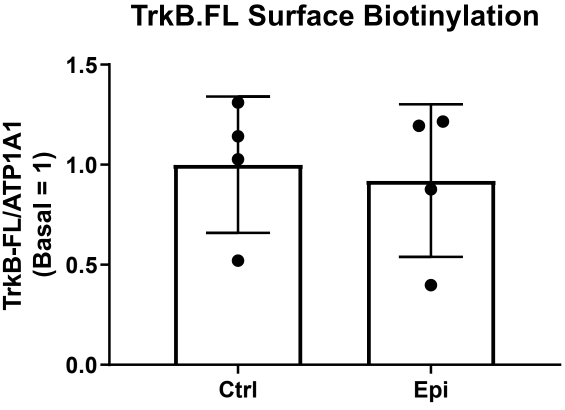
